# Analysis of clinical manifestations and risk factors of HIV-1 associated thrombocytopenia in a general teaching hospital in western China

**DOI:** 10.1101/2022.11.09.515900

**Authors:** Zhuoyun Tang, Zhonghao Wang, Tingting Wang, Jingyi Li, Chaonan Liu, Chuanmin Tao

**Author notes:** Correspondence to: Department of Laboratory Medicine, West China Hospital of Sichuan University, No. 37 Guoxue Lane, Chengdu 610041, China. E-mail address (CM Tao).

## Abstract

**Background:** Thrombocytopenia is one of the hematologic disorders that frequently accompany HIV-1 infection. The interaction of HIV-1 and platelets is crucial in the Blood-Brain Barrier’s (BBB) degeneration and causes neuroinflammation. The research aims to evaluate the prevalence and risk factors for HAT (HIV-1 associated thrombocytopenia) and summary the features of HAT-related neuroinflammation.

**Method:** A retrospective study including 221 HAT patients was conducted in West China Hospital from 2017 to 2021. Clinical and laboratory data was analyzed the prevalence and risk factors for HAT and HAT-related neuroinflammation.

**Results:** The prevalence of HAT was 11.06% and most patients were men (76.92%), elderly (≥50, 55.21%) and 63.80% patients were mild thrombocytopenia. CD4^+^ T cell count, platelet crit (PCT) and the rate of large platelets (P-LCR) were significantly different between HAT group and control group (*P* < 0.001, *P* 0.001, *P*=0.002). CD4^+^ T cell count <200 cell/μL (*P*=0.001) was an important risk factor in the occurrence of HAT while advanced age and high viral load were closely related to the occurrence of HAT. HAT-related neuroinflammation patients were mostly distributed in male (*X*^2^=10.066, *P*=0.007), with higher viral load (*X*^2^=12.297, *P*=0.006) and advanced age (*X*^2^=11.721, *P*=0.02) with neuropsychiatric symptoms and rising level of inflammatory factors like IL-6 and proteins in CSF.

**Conclusion:** HAT and HAT with neuroinflammation cannot be ignored in HIV-1 infection because of the activation of monocytes, macrophages and microglia, further causing thrombocytopenia and neuroinflammation. Advanced age, lower CD4^+^ T cell count and high viral load were closely related to the occurrence of HAT and HAT-related neuroinflammation.

## 1. Introduction

Human Immunodeficiency Virus-1 (HIV-1) disrupts human immunity, leading to cancers and other opportunistic infections, etc. Anemia, leukopenia, and thrombocytopenia are complications of hematologic disorders that frequently accompany HIV-1 infection and AIDS ^[1]^. The prevalence of HIV-1 associated thrombocytopenia (HAT), which can occur at any stage of the infection, ranges from 5.9% to 40%^[2–4]^. Thrombocytopenia may be the first clinical presentation in asymptomatic HIV-1-infected patients and may progress over time, leading to severe bleeding ^[5]^. The occurrence of HAT has not yet been determined. However, factors such as age, viral load, CD4^+^ T cell counts, and others may be related to the frequency of HAT. Additionally, the Blood-Brain Barrier (BBB) can be broken down by HIV-1 and its proteins, including Tat, allowing them to enter the central nervous system and cause neuroinflammation ^[6–7]^. The interaction of HIV-1 and platelets is crucial in the BBB’s degeneration, according to serval studies ^[8]^. Here, we integrated data from a large general teaching hospital in China with 4300 beds to evaluate the prevalence and risk factors for HAT and the features of HAT-related neuroinflammation.

## 2. Method

### 2.1 Study population

A retrospective study was conducted in West China Hospital with 4300 beds and a catchment population of approximately 16.33 million in Sichuan, China. The study included 221 HAT samples delivered to the laboratory, from inpatients and outpatients from 2017 to 2021. This study was reviewed by the ethics committee (approval number: 2020742).

### 2.2 Study-outcome definitions

HIV-1-infected patients were defined according to the guidelines for the diagnosis and treatment of HIV/AIDS in China (2021 edition). HAT patients were defined as HIV-1-infected patients with thrombocytopenia, which was defined as PLT count <100 × 10^9^/L. Mild thrombocytopenia was defined as a PLT count between 99 × 10^9^/L and 50 × 10^9^/L. Moderate thrombocytopenia was defined as a PLT count between 30 × 10^9^/L and 49 × 10^9^/L and severe thrombocytopenia as a PLT count <30 × 10^9^/L. HAT-related neuroinflammation was defined as HAT with neuroinflammation symptoms such as headache, dizziness and consciousness disorders, ect. HIV-1-infected individuals without thrombocytopenia were considered as control groups for viral load, CD4^+^ T cell counts, platelet crit (PCT), the rate of large platelets (P-LCR), and platelet distribution width (PDW). Quantities were paired 1:2 for each group. There were no significant differences in sex and age between the disease group and the control group (*P*>0.05).

### 2.3 Data collection

The following clinical and laboratory data was extracted from the electronic medical record system including sex, age, patient source, lymphocytes count, cut-off index (COI) of screening test, viral load, CD4^+^ T cell count, PCT, P-LCR and PDW. For HAT-related neuroinflammation, medical records were reviewed including protein in cerebrospinal fluid (CSF), IL-6 in serum and clinical diagnose besides the above indicators.

### 2.4 Statistical analysis

Data were recorded by Excel and analyzed by SPSS 23.0 and Origin 2022. Quantitative data were expressed as median and interquartile range (IQR). Noncategorical variables were assessed by t-test and categorical variables were counted by chi-square test or Fisher’s exact probability test. Univariate and multivariate logistic regression analysis were executed to assess the risk factors on the prevalence of HAT. And the outcomes were displayed as adjusted odds ratios (OR) with their respective 95% confidence intervals (CI). A P-value ≤ 0.05 was considered significant.

## 3. Results

### 3.1 The clinical and laboratory characteristics of HAT

A total of 1 998 samples were confirmed positive by Western Blot from 2017 to 2021 and 221 of them were finally diagnosed as HAT, with a prevalence of 11.06% (221/1 998). The prevalence steadily increased from 10.9% in 2017 to 18.67% in 2021. The clinical and laboratory characteristics of 221 HAT samples were summarized in Table 1. Most patients were predominantly male (76.92%), elderly (≥50, 55.21%), with high COI value, which the median COI value was 350.85 (173.30,724.25) and 63.80% patients were mild thrombocytopenia. Besides, the prevalence between patients under 50 years old and above 50 years old was totally different (*X*^2^=458.00, *P* < 0.001). Most HAT patients (71.95%) had low lymphocytes counts, indicating low CD4^+^ T cell counts.

**Table 1.**
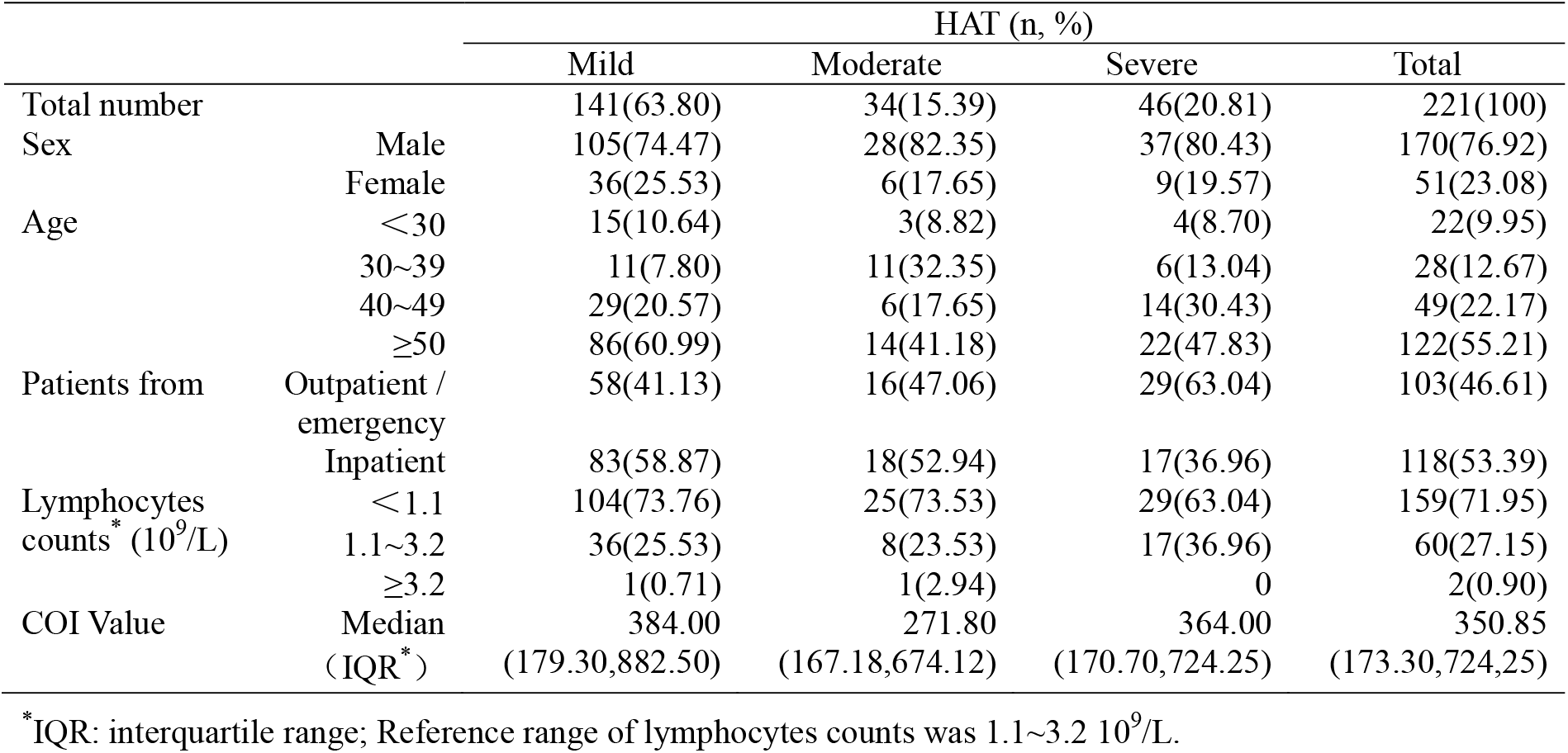
Clinical and laboratory characteristics of 221 HAT samples

### 3.2 Platelet parameters and risk factors of HAT

116 HAT patients underwent RNA testing, 120 HAT patients received CD4^+^ T cell count data, and 25 HAT patients, respectively, received PCT, P-LCR, and PDW results. The majority HAT patients (64.66%) had viral load ≥10^5^ cp/mL, 77.5% HAT patients had CD4^+^ T cell count < 200 cell/μL and 92% HAT patients had low PCT (<0.19). HAT patients had an average CD4^+^ T cell count of 123.11 cell/μL, which was substantially lower than that of the control group (*t*=3.565, *P* < 0.001). The values of PCT and P-LCR were significantly different between the HAT group and control group (*t*=5.372, *P* < 0.001; *t*=3.274, *P*=0.002). The viral load (*t*=1.540, *P*=0.125) and PDW values (*t*=1.150, *P*=0.254), on the other hand, showed no discernible variations. The variations between the HAT group and control group were showed in Figure 1.

**Figure 1.**
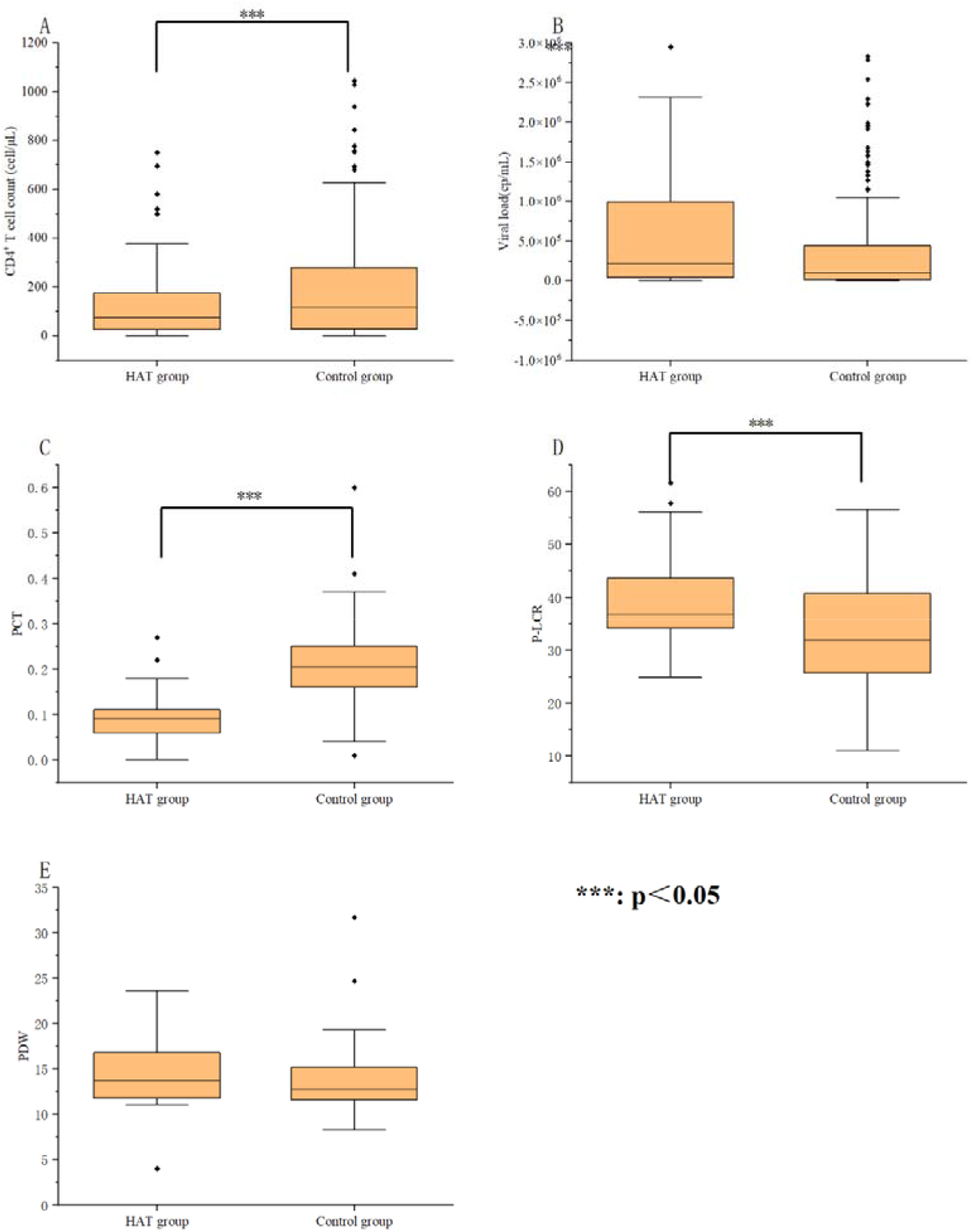
CD4^+^ T cell count, VL, platelets and related parameters of 221 HAT samples and control group

Regarding the frequency of HAT in the various groups, there were significant differences between CD4^+^ T cell count <200 cell/μL and ≥200 cell/μL (*X^2^*=248.00, *P*=0.019); viral load <10^5^ cps/mL and ≥10^5^ cps/mL (*X^2^*=119.00, *P* < 0.001) (Figure 2).

**Figure 2.**
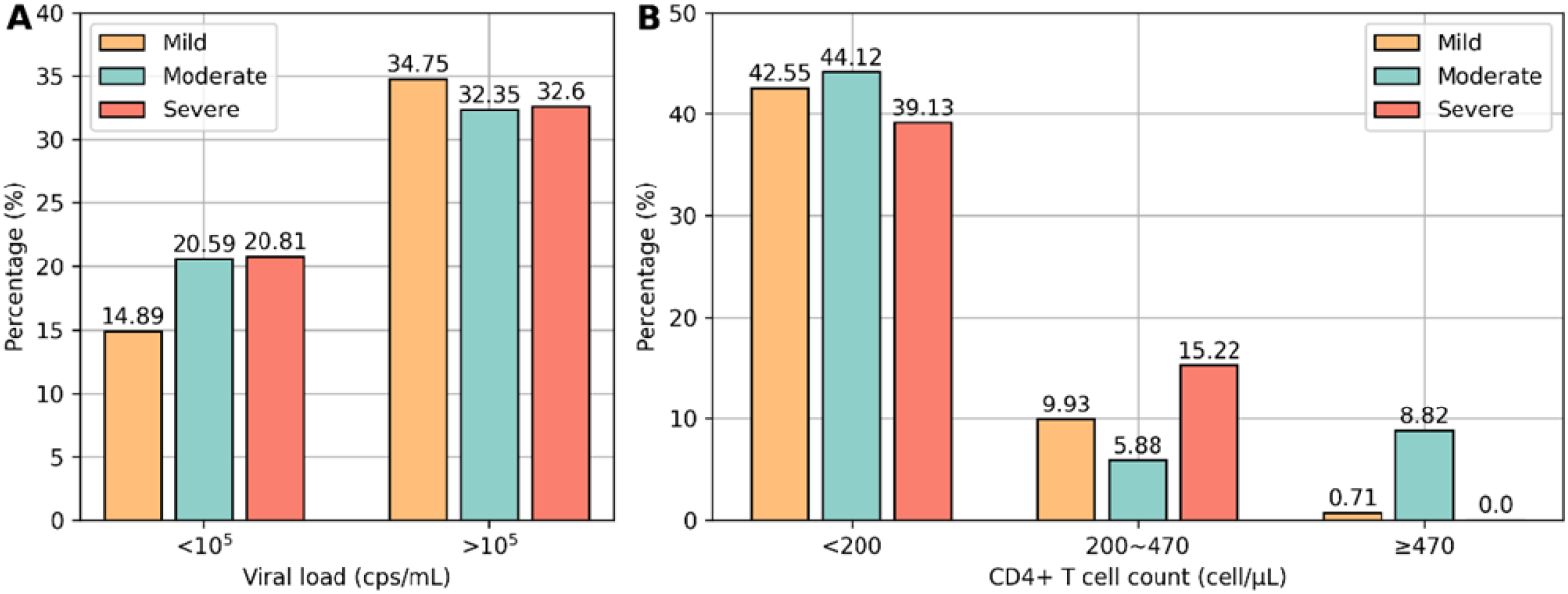
VL and CD4^+^ T cell count of mild, moderate, severe HAT samples

Univariate and multivariate logistic regression analysis both indicated CD4^+^ T cell count <200 cell/μL (OR=1.003, 95% CI=1.001-1.005, *P*=0.001) was risk factor in the occurrence of HAT. However, results showed high viral load (*P*=0.200) were closely related to the occurrence of HAT, although there was no significant difference, which was consistent with the results above. Most patients with advanced age (70.49%), high viral load (65.33%) and low CD4^+^ T cell count (64.52%) were distributed in the mild thrombocytopenia group.

### 3.3 The clinical and laboratory characteristics of HAT-related neuroinflammation

22 patients suffered from HAT-related neuroinflammation with a prevalence of 1.10% (22/1 998). Most patients were from emergency department (63.64%, 14/22). Neuropsychiatric symptoms accounted for a large proportion of newly diagnosed symptoms, including headache (27.27%,6/22), dizziness (22.73%,5/22), disturbance of consciousness (18.18%,4/22) and intracranial infection (18.18%,4/22). As for laboratory characteristics, 15 patients received the CSF biochemical testing and 13 patients had serum IL-6 results. 73.33% (11/15) patients showed high level of total trace protein exceeding normal and 84.62% (11/13) patients showed high level of IL-6 exceeding normal. Compared with normal HAT patients, patients with HAT-related neuroinflammation were mostly distributed in male (*X*^2^=10.066, *P*=0.007), with higher viral load (*X*^2^= 12.297, *P*=0.006) and advanced age (*X*^2^=11.721, *P*=0.02). Although there was no significant difference in CD4^+^ T cell count, the result indicated a lower CD4^+^ T cell count in patients with HAT-related neuroinflammation. Table 2 and Figure 3 compared patients with HAT-related neuroinflammation and without neuroinflammation.

**Figure 3.**
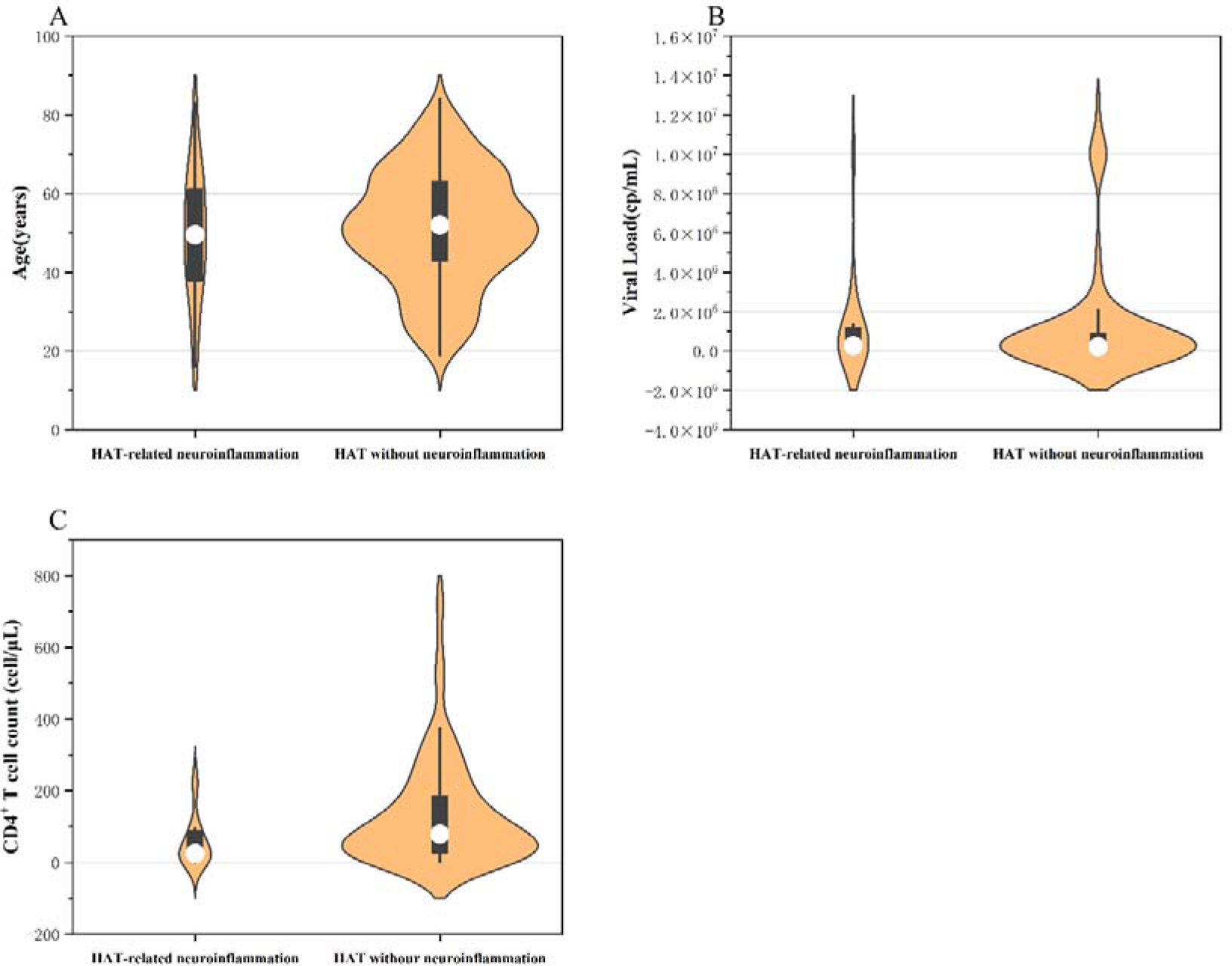
Comparison of HAT-related neuroinflammation and without neuroinflammation (Age, viral load and CD4^+^ T cell count)

**Table 2.**
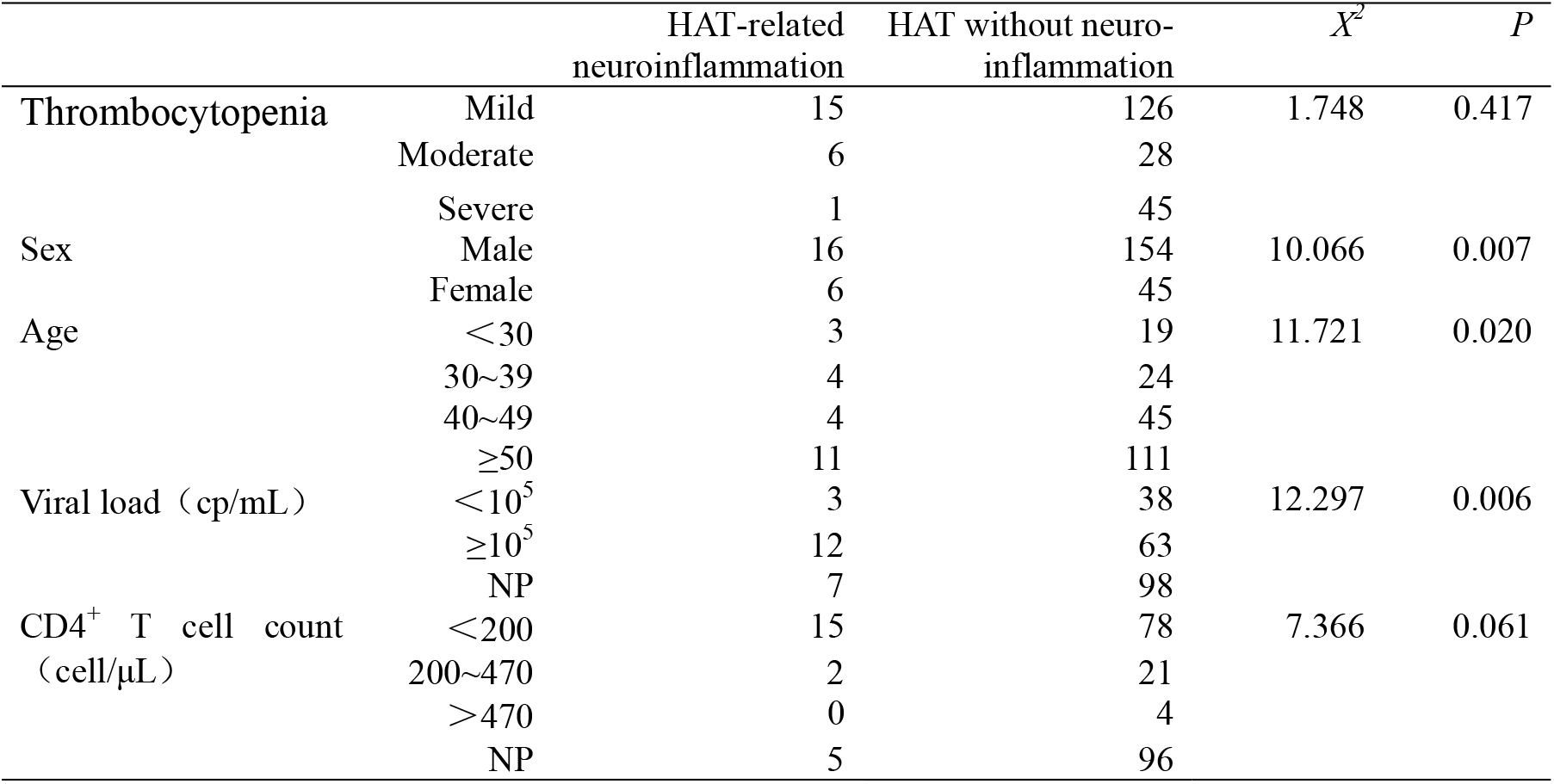
Comparison of HAT-related neuroinflammation and without neuroinflammation

## 4. Discussion

Sichuan province has one of the highest rates of HIV-1 infection in China and a high incidence of thrombocytopenia ^[9–10]^. Thrombocytopenia, a common hematological disease, may be the first clinical sign in HIV-1-infected patients. The study showed the prevalence of HAT was 11.06%, which contrasted with results in other publications that ranged from 10% to 30%^[11–16]^. The prevalence was similar with HIV-1-associated tumor, HIV-1-associated fungal infection, but less than HIV-1-associated tuberculosis ^[17–20]^. The findings indicated that thrombocytopenia occurred more frequently in HIV-1-infected patients, especially before the initiation of antiretroviral therapy, compared to the general population ^[1]^. The majority of HAT patients were male, elderly, and with high COI. In terms of the majority of HAT, males constitute the category with the highest prevalence of HIV-1 infection ^[21–22]^. Elder patients had more severe illnesses and underlying problems, which contributed to a few HIV-1 infection consequences, including HAT ^[23]^. High COI levels probably suggest an early stage of infection and rapid virus replication, which resulted in aberrant megakaryocyte differentiation, viral-induced suppression of hematopoietic stem cell proliferation, and cytokine dysregulation ^[24]^.

The results showed that CD4^+^ T cell count, PCT and P-LCR were significantly different between HAT group and control group. PLT, PCT, P-LCR, and PDW, respectively, represent the quantity, the size of platelets and a measure of the product’s degree of variation and the percentage of large platelets, are considered to be the markers of platelet activation ^[25]^. PCT can represent the degree of platelet activation, and a simultaneous decrease in PLT and PCT indicates excessive consumption of platelets ^[26]^. Additionally, high P-LCR levels indicate an increase in newborn or relatively immature platelets with large volumes, more densed particles, and robust activity that reacts vigorously to inflammatory mediators or pro-inflammatory chemokines ^[27]^. The findings therefore demonstrated that PLT and PCT decreased after viral infection, and the feedback increased platelet activation and production, resulting in the increase of P-LCR.

As our study observed, a significant low CD4^+^ T cell count (< 200 cell/μL) was found to be the driving factors of HAT. Lower CD4^+^ T cell counts have been linked to an increased risk of thrombocytopenia in other studies ^[28–30]^. A study discovered that among homosexual males, a fall in PLT quantity indicated a sharp decline in CD4^+^ T cell counts ^[3]^. Besides, advanced age and high viral load were closely related to the occurrence of HAT which were similar with previous studies ^[28–30]^. High viral load suggested active viral replication, while low CD4^+^ T cell count signaled the immune system’s demise. When platelets and blood cells start to decline, it indicates that the patient’s condition is rapidly deteriorating and worsening, which increases the risk of infection with bacteria, fungi, and viruses, among other things.

On the other hand, this study focused on the HAT-related neuroinflammation, which the clinical symptoms were mainly headache, dizziness and disturbance of consciousness, etc. HIV-1-associated neurocognitive disorders (HAND) are one of the most important complications as an end-organ manifestation of HIV-1 infection. Neuroinflammation is a precursor phenotype of HAND and is difficult to diagnose, which deserves attention. Compared to HAT without neuroinflammation, these patients shared several traits, such as advanced age and higher viral load. Additionally, it revealed a pattern where patients with HAT and neuroinflammation had lower CD4^+^ T cell counts and the increasing levels of protein in CSF and IL-6 in serum. The elevation of protein in CSF is a serious warning, especially in light of that some researches showed elevated β_2_-microglobulin in CSF was an important predictors of AIDS related dementia ^[31–32]^. According to several research, HIV-1 cannot directly harm neurons but can infect peripheral blood monocytes, pass through the BBB, enter brain tissue. Additionally, platelets can influence the BBB’s integrity and permeability in vitro, worsening the BBB’s breakdown ^[33–36]^. Subsequently, HIV-1 expresses neurotoxic compounds like gp120, Tat, Nef proteins to enhance the neurotoxicity and cause neuroinflammation, leading to the increase of inflammatory factors like TNF-α, IL-6, etc ^[35,37–40]^. Furthermore, one of mechanism that HIV-1 exerting neurotoxicity is the interaction between HIV-1 and the monocyte-platelet aggregates, which results in aberrant platelet activation and thrombocytopenia ^[41–42]^.

Our study has some limitations owing to its retrospective design. First, research from single center cannot provide a more accurate representation of the epidemiology of HAT and HAT with neuroinflammation. Second, some data such as viral load, CD4^+^ T cell count and platelet parameters was partly missing.

In conclusion, HAT and HAT with neuroinflammation cannot be ignored in HIV-1 infection. Thrombocytopenia can be caused by abnormally platelet activation and resulting in the decrease of PCT and the increase of P-LCR. High levels of inflammatory agents and proteins induced by HIV-1’s neurotoxicity resulted in neuroinflammation-related symptoms and a worse prognosis. Currently, there is no definitive diagnosis for HAT and HAT with neuroinflammation, therefore this research would certainly give some inspiration on how the condition should be treated. Additionally, the diagnostic methods and pathogenesis remain to be further studied.

## Authors Contributions

Chuanmin Tao conceived the initial concept, reviewed and edited the manuscript. Zhuoyun Tang wrote the original draft. Zhuoyun Tang and Chuanmin Tao designed the study. Zhuoyun Tang, Jingyi Li and Chaonan Liu performed the experiments and did the formal analysis. Zhonghao Wang and Tingting Wang revised the original draft of manuscript.

## Acknowledgements

This work was supported by the National Natural Science Foundation of China (82102485, Z.W.).

